# Single Cell Viewer (SCV): An interactive visualization data portal for single cell RNA sequence data

**DOI:** 10.1101/664789

**Authors:** Shuoguo Wang, Constance Brett, Mohan Bolisetty, Ryan Golhar, Isaac Neuhaus, Kandasamy Ravi

**Affiliations:** Bristol-Myers Squibb, 311 Pennington Rocky-Hill Rd, Pennington, NJ 08534

## Abstract

**Motivation:** Thanks to technological advances made in the last few years, we are now able to study transcriptomes from thousands of single cells. These have been applied widely to study various aspects of Biology. Nevertheless, comprehending and inferring meaningful biological insights from these large datasets is still a challenge. Although tools are being developed to deal with the data complexity and data volume, we do not have yet an effective visualizations and comparative analysis tools to realize the full value of these datasets.

**Results:** In order to address this gap, we implemented a single cell data visualization portal called Single Cell Viewer (SCV). SCV is an R shiny application that offers users rich visualization and exploratory data analysis options for single cell datasets.

**Availability:** Source code for the application is available online at GitHub (http://www.github.com/neuhausi/single-cell-viewer) and there is a hosted exploration application using the same example dataset as this publication at http://periscopeapps.org/scv_tirosh.

## 1 Introduction

Single cell RNA sequencing allows the study of RNA expression from individual cells. This enables studying expression differences at single cell resolution and identifying transient cell states and identity. It can reveal aspects of biology that are not possible with bulk RNA sequencing such as uncovering trajectories of cell type lineages during development (Farrell et al., 2018) or existence of intermediate and rare cell types (Villani et al., 2017). There has been an exponential growth in the number of single-cell RNA-sequencing datasets in the past few years, thanks to advances made in technology, especially in nanodroplet- and microfluidics-based systems. Measuring large numbers of single cells, in the millions, has become feasible and affordable (Svensson et al., 2018).

Due to rapid advancement of single cell technology, there has been increase in data volume and complexity, bringing with it many new computational challenges. With the increasing availability of many large data sets such as Human Cell Atlas Data (Regev et al., 2017) deriving biological insights has become a critical need. It requires effective ways of visualizing the data and the ability to perform exploratory data analysis (portals.broadinstitute.org/single_cell; Zhang et al., 2018). Currently available tools are either missing the scalability to host multiple data sets or provide limited visualization options with minimal or no exploratory analytical capabilities.

We designed an exploratory tool, single cell viewer (SCV) which is an R shiny application for visualization and analysis of single cell data. It addresses the following needs: 1) Interactive rich visualization, 2) generating high quality figures suitable for publication, 3) provide exploratory analysis including differential gene expression analysis between clusters or selected set of cells. We implemented SCV using open source computing infrastructure such as periscope (Brett et. al, 2019) and canvasXpress (Neuhaus et. al, 2019).

## 2 Methods

SCV data visualization requires a single .rds file, which is an S4 class object defined by the R toolkit Seurat (Butler et al., 2018). We suggest creating a one R object of data aggregated from many patients or samples in an experiment. To be compatible with SCV, a Seurat object must have the following slots: 1) obj@assays slot holds the raw expression matrix that is set as the; 2) obj@reduction slot holds the “tsne” slot containing the tSNE coordinates of each cell; 3) obj@meta.data contains information associated with each cell such as treatment, responder status, or other information such as nCount_RNA, nFeature_RNA etc.; 4), which defines cell cluster identity, needs to be set to one of the identity variable in the obj@meta.data; 5) obj@misc with pre-computed top 10 differentially expressed genes (DEG) stored in table format at obj@misc$DE$top10, which will be displayed on the summary page (see “Visualization Design”). A larger list containing top 30 DEG are stored at obj@misc$DE$top30. Users also need to provide associated metadata about cells stored as a named list at obj@misc$meta.info, as shown in the provided template (see supplementary) and script to add information to the Seurat objects. Additionally, user needs to define a set of meta data at obj@misc$DataSegregate that are also present in obj@meta.data for advanced data filtering and segregation (see “Supplementary Methods” for more details).

There are two methods to obtain a compatible .rds file for SCV. The first is to start with an expression matrix for example .csv or .mtx files from CellRanger output (raw data matrix from multiple samples can be merged using Seurat or cellranger.aggr function) and run the standard Seurat analysis workflow (https://satijalab.org/, see “Supplementary Methods” for more details and example scripts). The Seurat workflow includes several steps. The analytical results from each step will be stored in slots, and users need to perform differential gene expression analysis separately to obtain the top10 and top30 DEGs and add required Meta information to be object.

Alternatively, users can also take advantage of pre-analyzed single cell RNA-seq datasets from publications. As long as the objects are created with slots as described above.

For comparative analysis from different patients or treatment conditions, corresponding metadata must be present in the object at obj@meta.data slot which can be loaded by built-in Seurat functions. SCV offers two-options for data filtering and segregation: 1) sub-setting with pre-defined Meta data information such as tissue type, response status, treatment, etc; and 2) further segregate the filtered data from step 1 into faceted panel charts using meta data. This feature enables users to select and perform analysis between subsets for example when computing differentially expressed genes.

## 3 Visualization Design

SCV provides accessible representations of the underlying data through a variety of rich visualizations. The initial plot on entry to the application in the Summary tab is a tSNE plot of the entire dataset. This plot shows all the cell populations and is colored by cluster. This is paired with a searchable, sortable, and downloadable summary table that lists differentially expressed genes by cluster including adjusted p-values and relative fold-change for either the top ten or thirty differentially expressed genes.

**Fig. 3.1.**
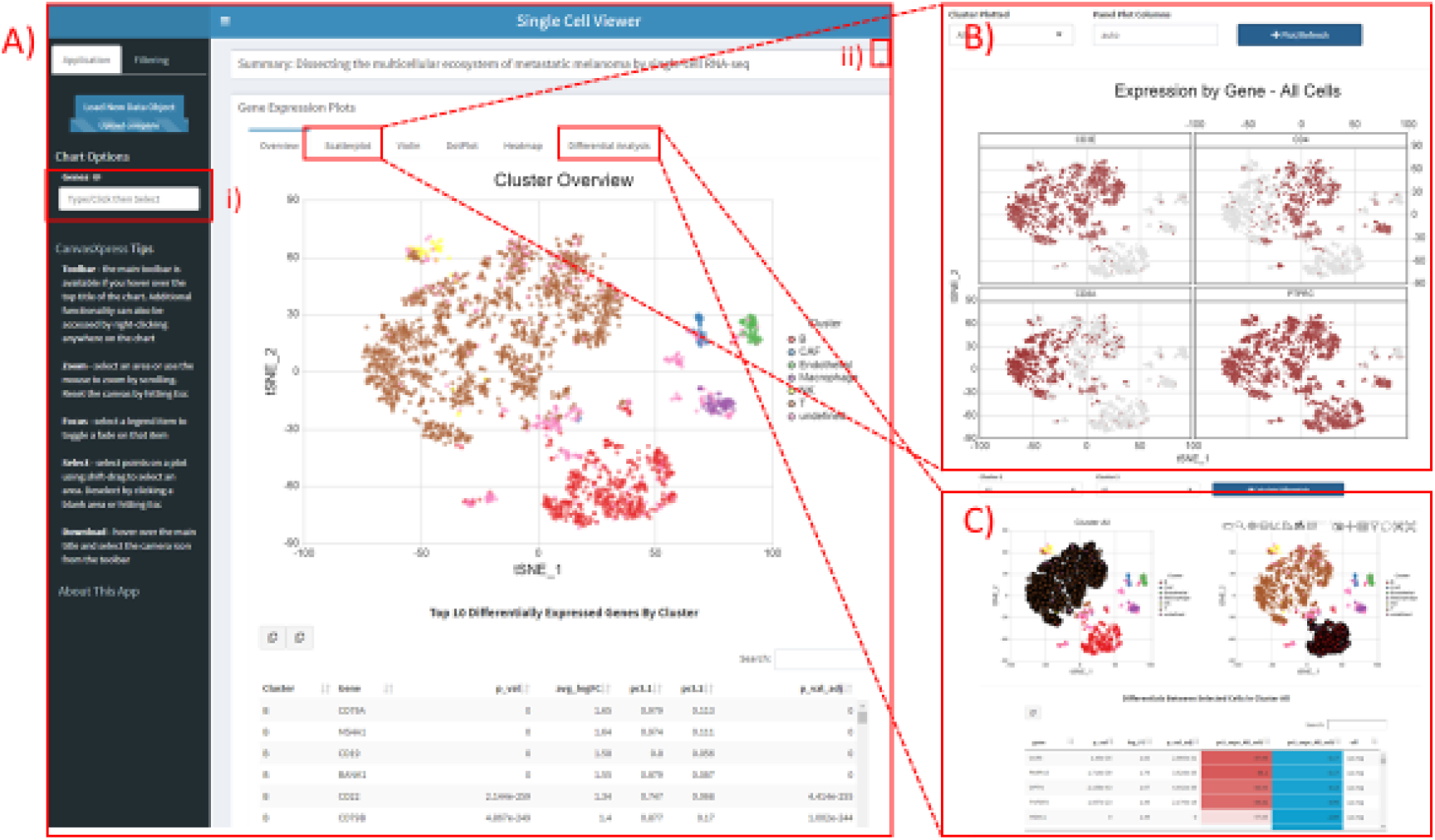
Visualization design of SCV. Showing on left is the overview tab (A), right are the scatter plot (B) and differential analysis (C) tabs.

Users can select genes of interest to further explore the dataset. Expression-level tSNE scatterplot as well as violin distribution plots with a boxplot insert allowing exploration of summary and detailed information by gene. To explore more than a few specific genes there are heatmaps and dot plots. These plots also have the option to include the top differentially expressed genes for comparison with any specific genes of interest. The dot plot showcases the number of expressing cells as well as the average expression value of selected genes for each cluster. The heatmap utilizes normalized values to reflect the original expression levels of the cells and clusters.

To explore beyond pre-computed differential expression values the application provides for a custom differential analysis to be performed. The user can select any two clusters or any two groups of cells and perform a differential expression of genes (DEG) analysis of those two groups on demand in the application. To ensure a performant DEG analysis there is a sorting and filtering step done where the application pre-selects up to 1000 genes for differential gene analysis. Most DEG analyses using the application take less than 30 seconds to complete. Computed DEG values are also provided to the user in a searchable, sortable, downloadable table including the p-value (Wilcoxon Rank Sum test), adjusted p-value (Bonferroni correction), log2 fold changes, percentage of cells expressing, and the effect (up-or down-regulated).

All charts in the application are implemented using canavasXpress, a highly functional JavaScript library (Neuhaus, et. al, 2019) to support data segregation, data filtering, extensive tooltips, zoom/pan, download, and many other functionalities.

## 4 Example dataset

For Demonstration of the SCV tool as an example we used published melanoma single cell RNA seq dataset (Tirosh et al., Science, 2016) which can be downloaded from https://portals.broadinstitute.org/single_cell/study/SCP11/melanoma-intra-tumor-heterogeneity. Once downloaded the first step is creating a SCV compatible R object using instructions as described in Supplementary Script. The object can be uploaded in to SCV by using upload button in the SCV application.

### 4.1 Overview Page

Once the object is loaded in SCV the first page or the overview page provides a summary of the combined object with annotated clusters. The overview page contains sidebar on the left, with two tabs. The first tab allows the users to create sub-plots on relevant tabs with customized gene list. The second tab enable users to perform global filtering of cells such as specific sub-population or cluster of interest.

On the right top, users can expand the summary box to view the metadata information associated with the object that was defined at obj@. Below the summary box is a summary tSNE plot showing all the cells and clusters.

The pre-computed top 10 differentially expressed genes for each cluster are displayed by default at the bottom as a table. The genes are first ordered by cluster then by average log fold change (ave_logFC). The un-adjusted p-value (p_val) and Bonfferoni corrected p-value (p_val_adj) indicates the significance level of the differential expression. We also display the percentage of cells that have non-zero expression of the gene for both the test cluster (pct.1) and the rest of the cells other than the test cluster (pct.2). The table can be downloaded as csv for both the top10 and top30 differential genes. The downloaded csv will match the format shown, which also matches the data frame saved at obj@misc$DE$top10.

**Figure 4.1.**
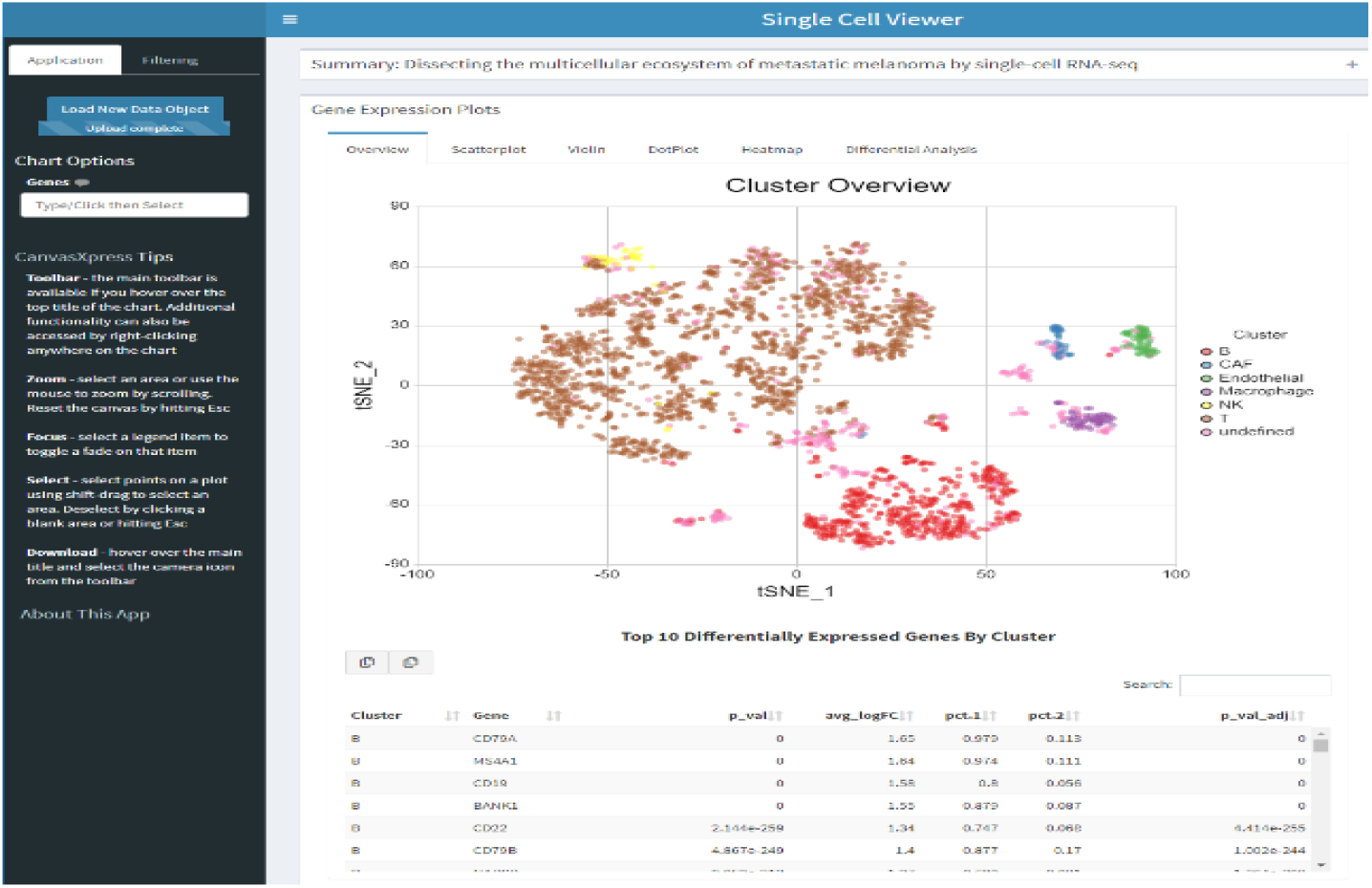
Overview Page showing cluster identity and differential gene list.

### 4.2. Scatter plot

Scatter plot or feature plot is a popular and intuitive way of looking at the expression of list of genes expressed in the object either from user provided list in the search box from the sidebar for all the cells in object of specific clusters. One can also set columns for displaying scatter plot.

**Figure 4.2.**
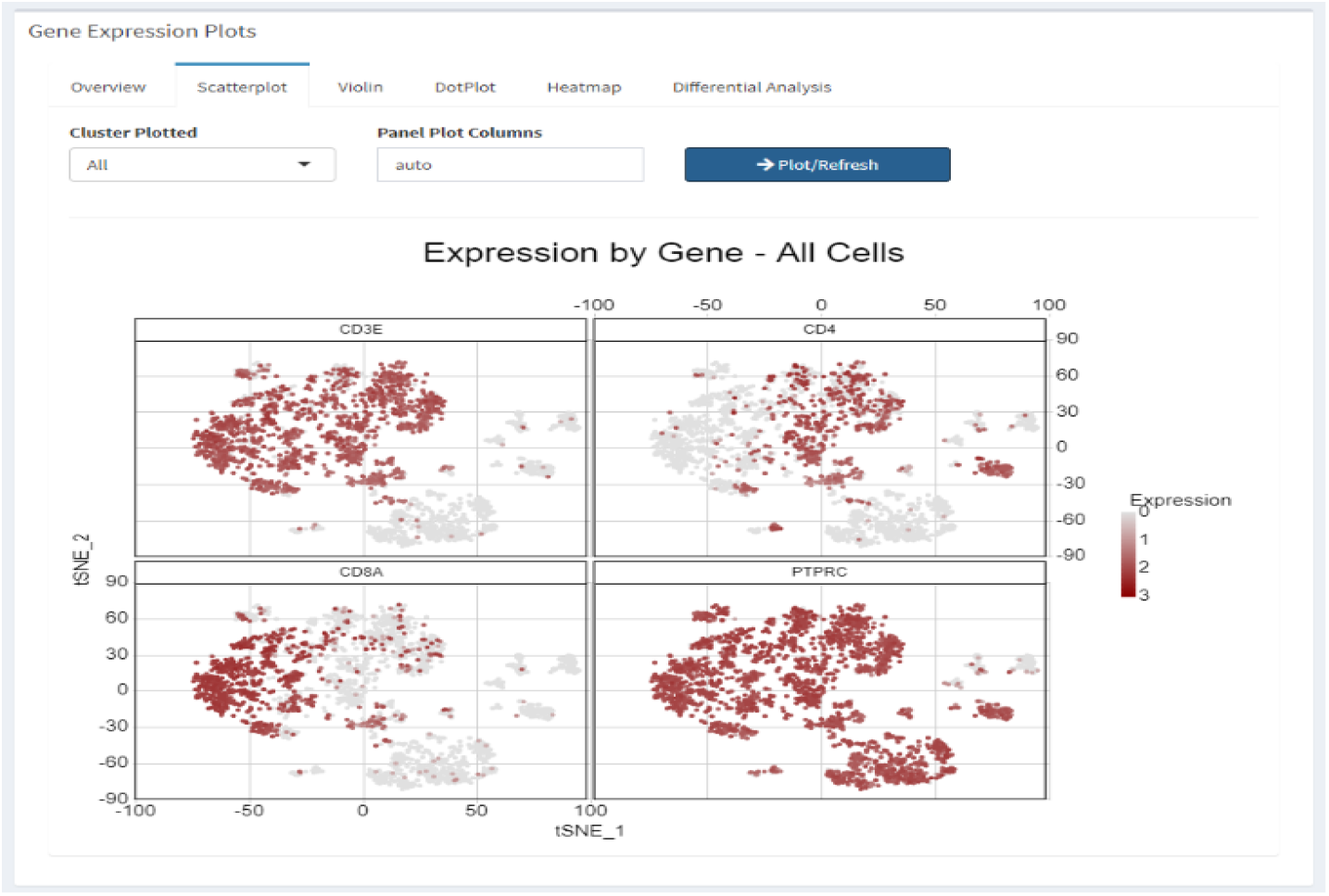
Scatter Plot showing expression of genes in individual cells.

### 4.3. Violin plot

Violin plots breaks down the expression summary for genes (selected in the sidebar) by clusters. A boxplot is overlaid, and users can hover over the distributions to show the actual values. The actual expression values of individual cells are hidden from the violin plot unless it is above 3rd quantile or below 1st quantile values (e.g. outliers).

**Figure 4.3.**
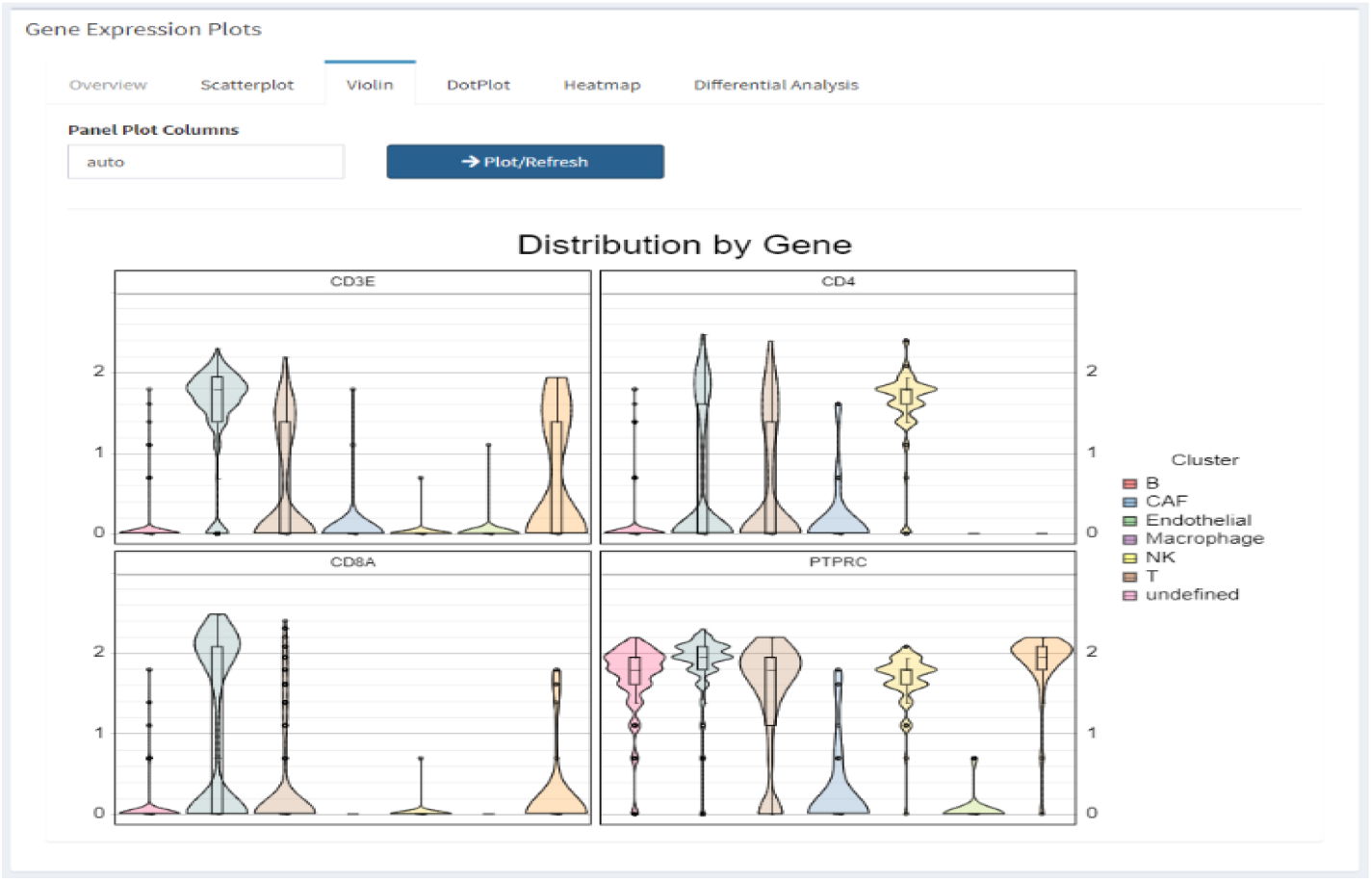
Violin Plot showing summary information for multiple genes in each cluster.

### 4.4. Dot Plot

The Dot Plot shows summary information similar to violin plot but condenses information for larger query gene sets (selected in the sidebar, and optionally adding the top10 or top30 genes). The color of the dots shows the average expression value and the size of the dots shows the percentage (%) of cells in each immune population that express the gene. The average expression levels are scaled the same way the Seurat package (Butler et.al., 2018) scales data for dot plots.

**Figure 4.4.**
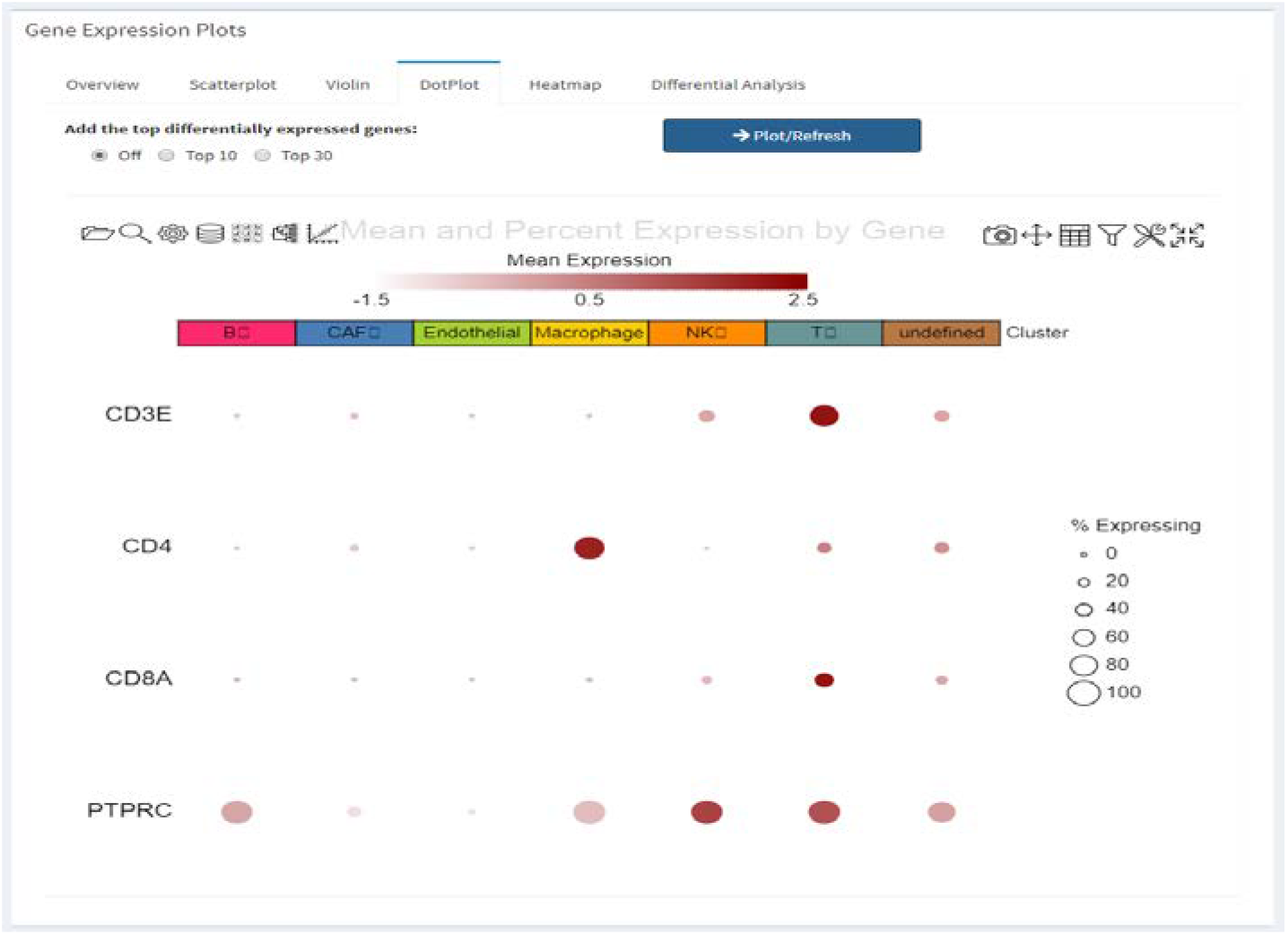
Dot Plot showing summary information for genes in each cluster.

### 4.5. Heatmap Plot

The Heatmap shows expression values of larger query gene sets for individual cells grouped by clusters. The color showings the value from the active assay and is great way to diagnose the clustering accuracy and identify potential sub-structures.

**Figure 4.5.**
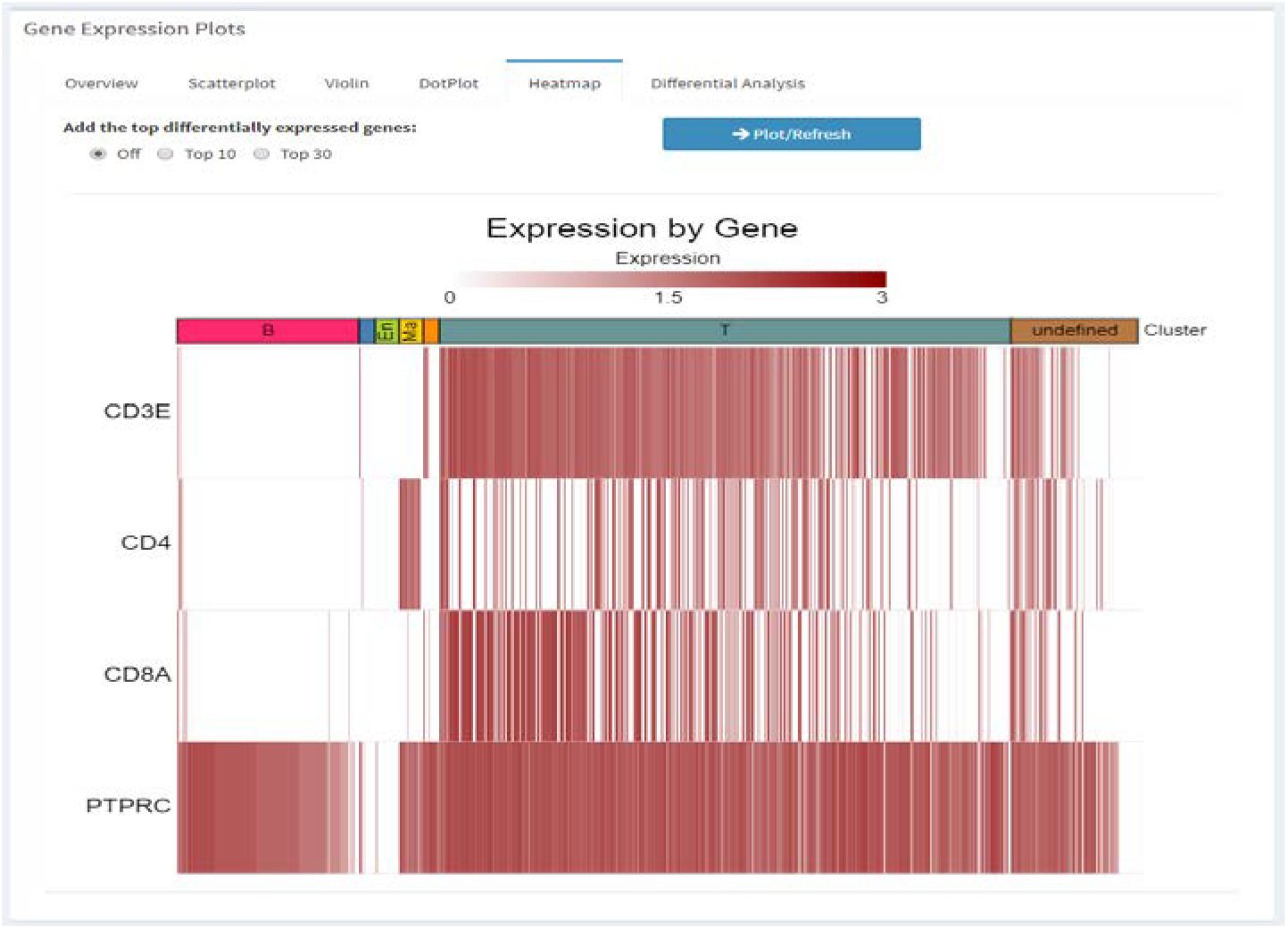
Heatmap showing expression (raw count) for genes in each cluster by individual genes.

### 4.6. Differential Analysis

One of the most powerful features that SCV provides is the ability to perform differential gene analysis. Users can select two different clusters to perform cluster DE analysis, or if the same cluster is selected users can select two groups of individual values on either chart, using the rectangular selection or lasso selection of arbitrary shapes, and perform sub-cluster DE analysis. Differential Expression gene analysis is performed using the Wilcoxon Rank Sum test with the results shown below the plot in a table. In the table pct_expr_All_[cluster1|set1] and pct_expr_All_[cluster2|set2] are color-coded to reflect the percentage of cells expressing the gene for cluster1/set1 and cluster2/set2, respectively. The user can also download the DE data as a csv/tsv file by clicking the 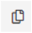 button.

**Figure 4.6.**
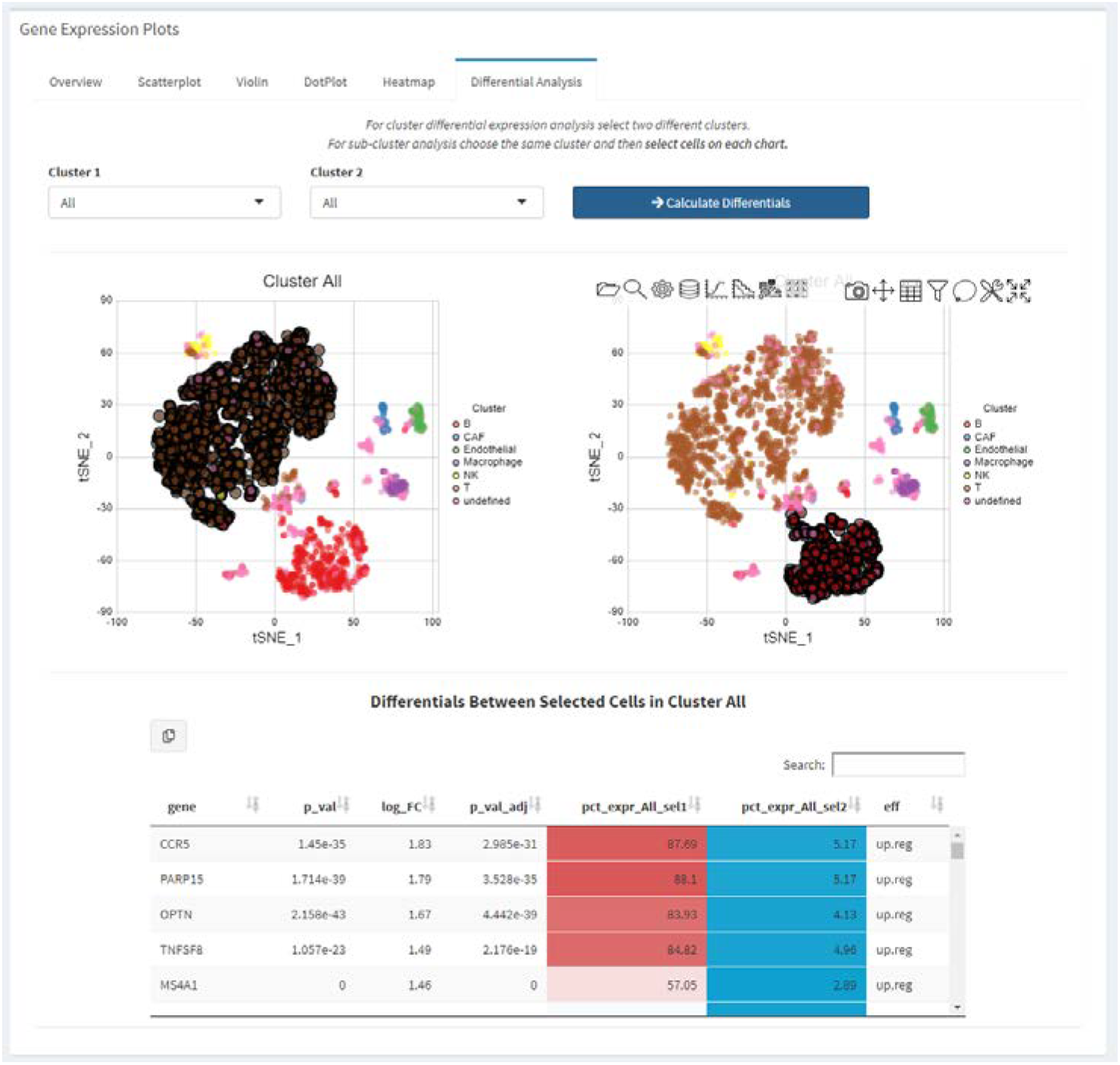
Differential Analysis with in SCV.

### 4.7. Advanced data sub-setting and segregation

Advanced data sub-setting and segregation is another important feature built-in to SCV to facilitate complex data partitioning and analysis of the combined object. It allows users to perform comparisons like pre- and post-treatment, responder vs. non-responder, and check biological variations among patients. Below we demo how users can use data from a subset of patient aggregates, and perform differential gene analysis by comparing only the T cells from two different patients.

First, users can perform global data sub-setting on the second tab of the left panel (Figure 4.7.1). Filtering performed here will be reflected for all the visualizations and analysis performed subsequently and in real-time on the overview tSNE plot. For demo purposes, we arbitrarily unchecked the first 6 patients from the dataset, and the tSNE plot will reflects the updated changes in cells in real-time as the patients are unchecked.

**Figure 4.7.1.**
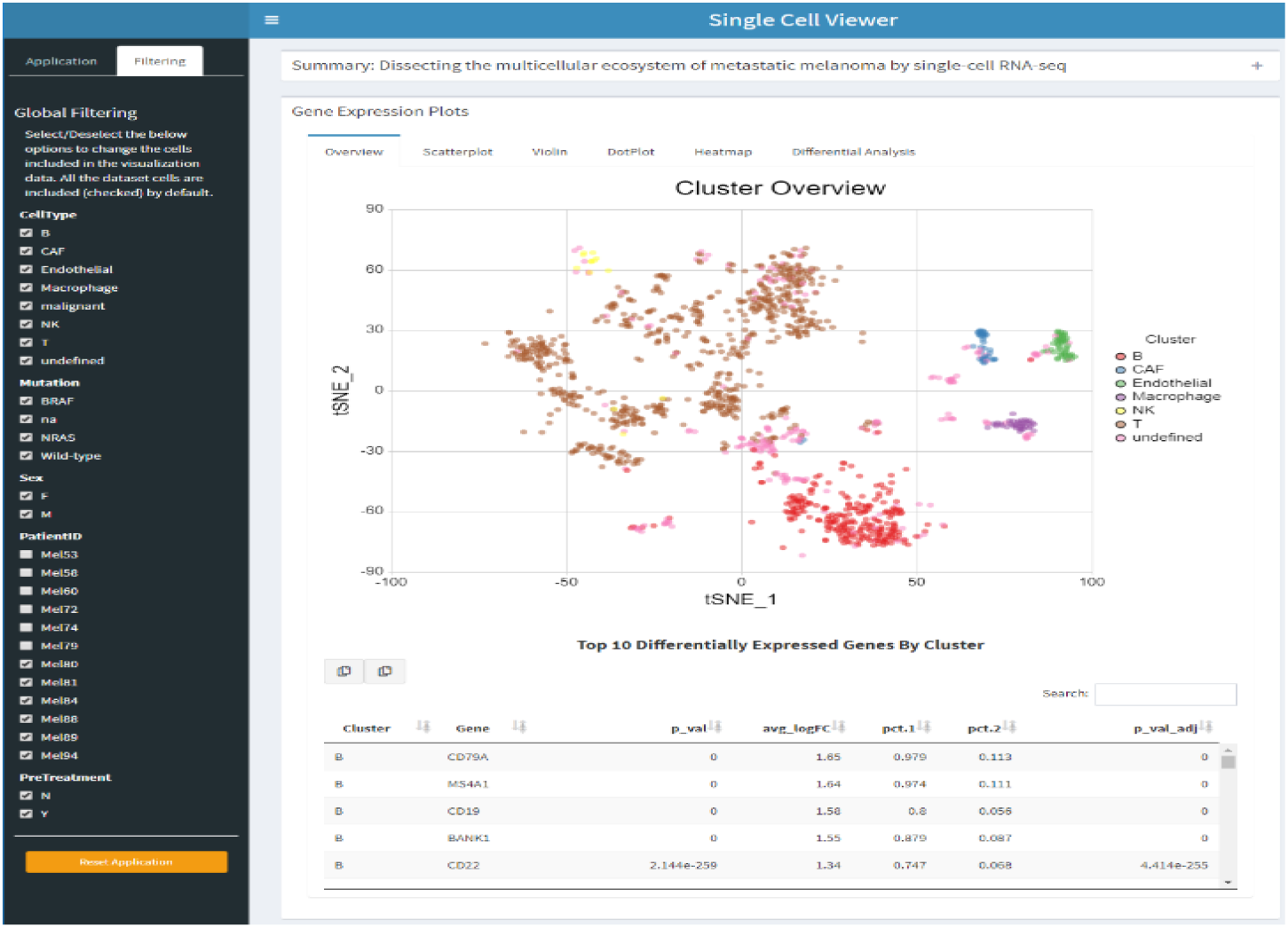
Data filtering based on meta information.

After de-selection, a scatterplot can be created of the same four genes as above showing expression in this particular subset of cells for only the last 6 patients (Figure 4.7.2).

**Figure 4.7.2.**
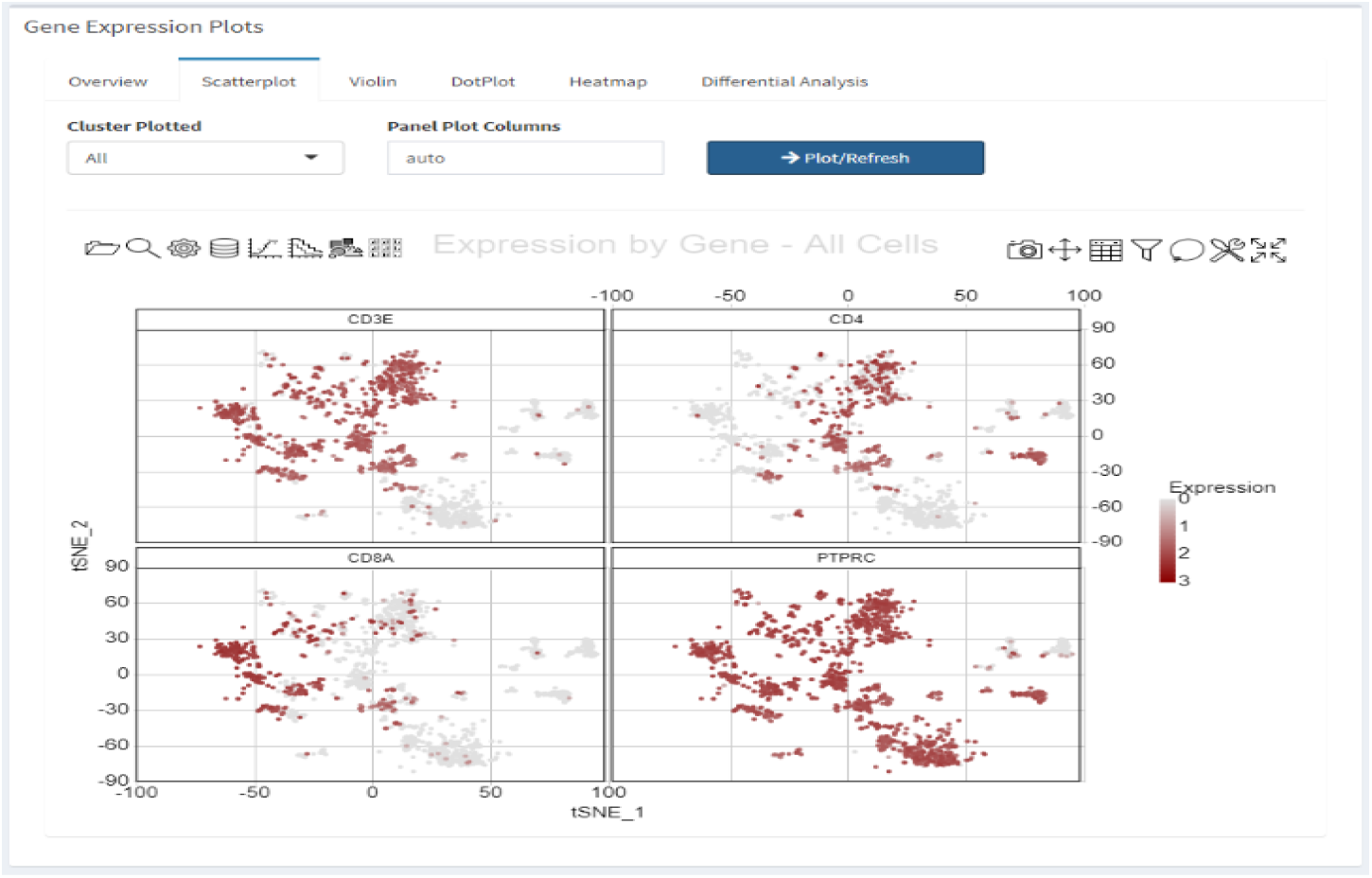
Scatter Plot showing gene expression that after data filtering which imposed as global effects across SCV.

Users can further segregate the cells on the plots into different sub-populations based on any one of the variables defined in the obj@misc@DataSegregation slot by clicking on the icon in the plot (Figure 4.7.3). For the demo data we provided the several metadata options for cell segregation including patients, mutation status, previous treatment history, etc. Below we segregate the cells further by patients to show the data filtering.

**Figure 4.7.3.**
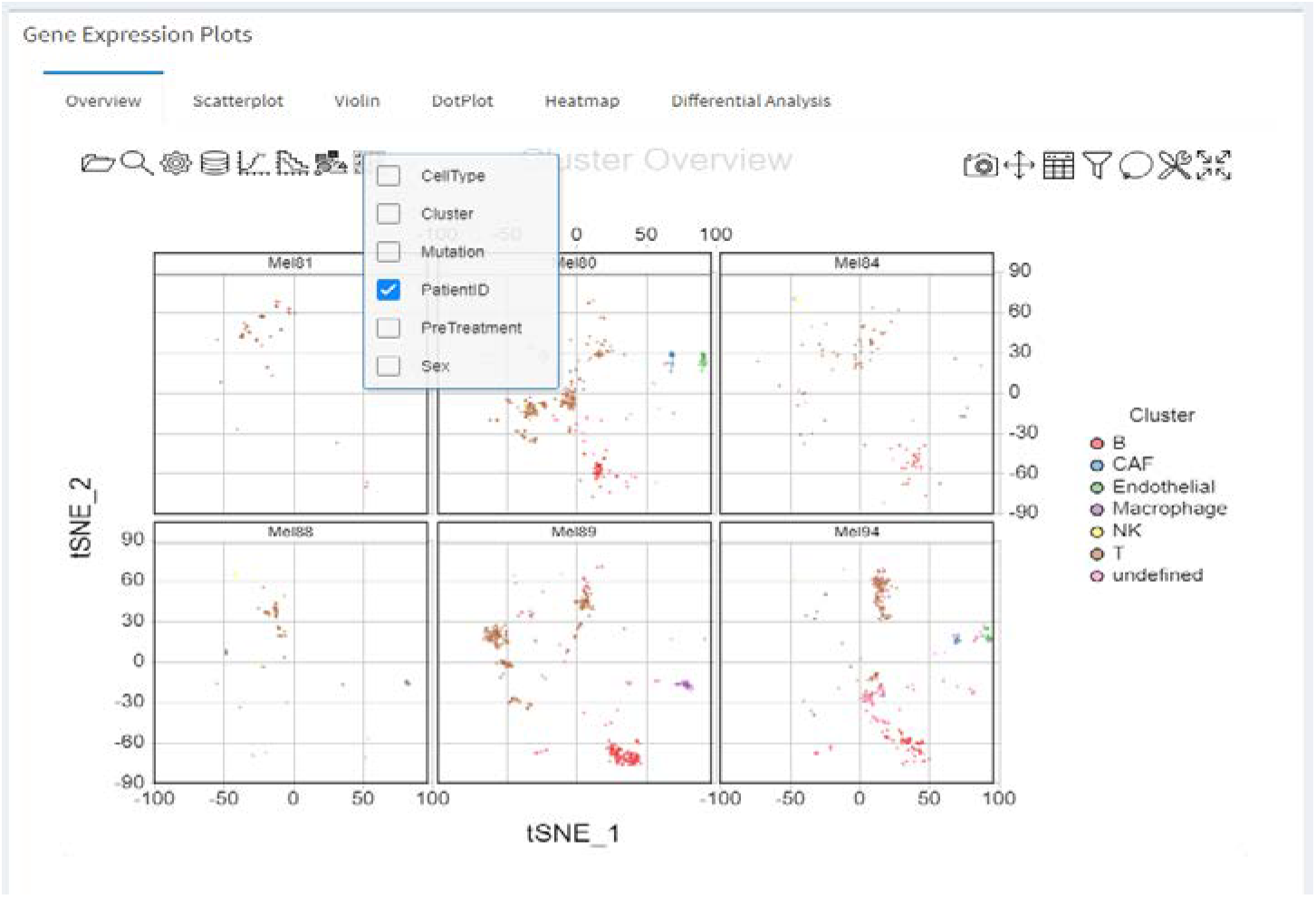
Data segregation using meta information, e.g. by Patient ID.

Next, in the “Differential Analysis” tab, the cells are segregated by patients in both plots (Figure 4.7.4) and T cells from patient Mel89 were selected as group 1 and T cells from patient Mel80 were selected as group 2. Differential analysis is then performed as previously described and the results here are between T cells only from these two patients.

**Figure 4.7.4.**
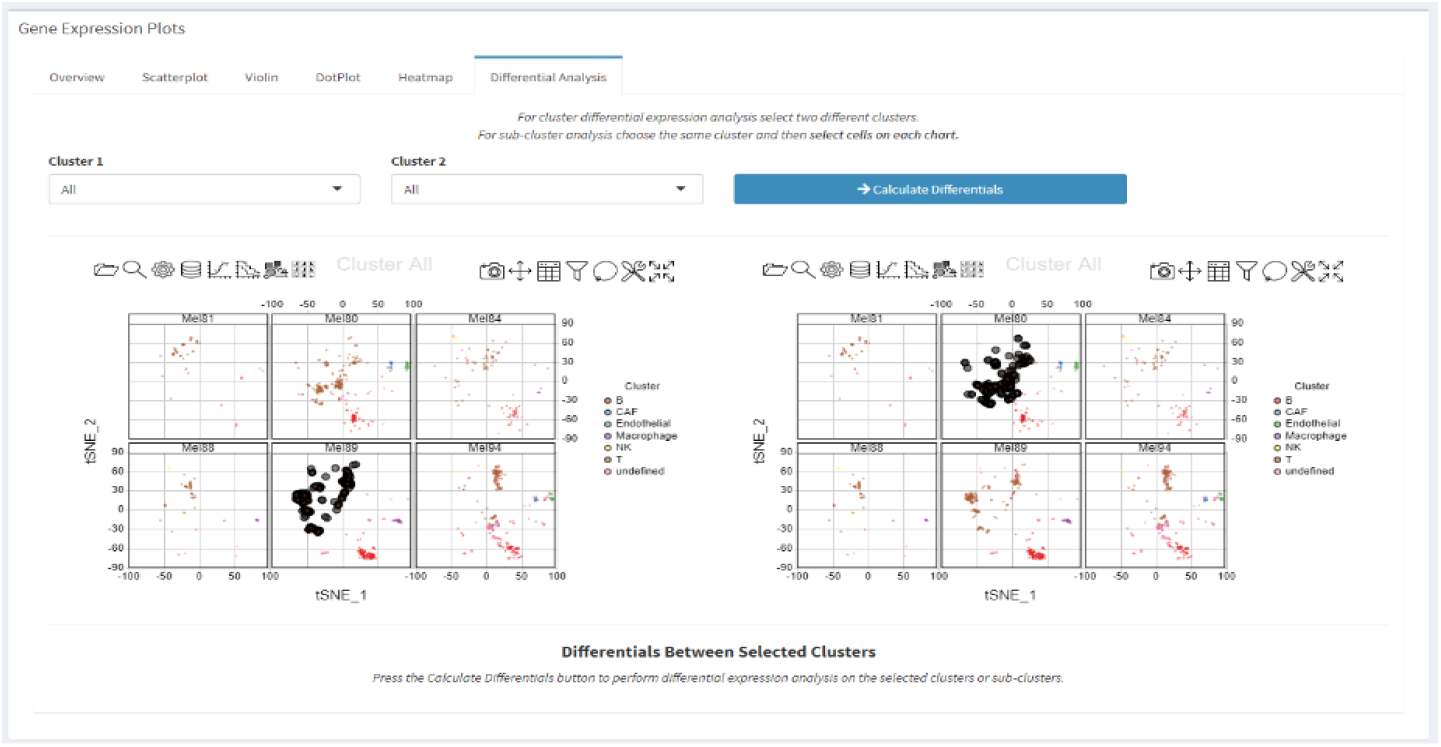
Differential gene analysis for T cells between two patients were enabled by filtering and segregating data.

## Discussion

The data import functionality is extensible to other formats such as h5 or loom format, although not currently supported. The application is easily updatable to load data from locations such as a local or remote file.

## Supporting information

SCV_Supplementary

