## Supplementary material for "Single Cell Viewer (SCV): An interactive visualization data portal for single cell RNA sequence data": SCV_Supplementary

### Supplementary Methods

#### 1. Demonstration dataset

For demo purposes we have used published melanoma dataset (Tirosh et al., Science, 2016), which can be downloaded from https://portals.broadinstitute.org/single_cell/study/SCP11/melanoma-intra-tumor-heterogeneity) and provided an example script ("supplementary script") to produce a Seurat version 3 object compatible with the single cell viewer.

#### 2. Single Cell Viewer (SCV) compatible Seurat Object Structure

SCV data portal visualization takes a single rds file, which is an S4 class object defined by the R toolkit Seurat (Butler et al., 2018, version 3) with multiple pre-defined slots to store layers of information. There are five slots that are required for the Seurat objects to be compatible with SCV.

1) the object required at least one assay matrix been stored and defined as the "active assay".

```
> myproject@assays$RNA[1:2,1:2]
2 x 2 sparse Matrix of class "dgCMatrix"
         Cy72_CD45_H02_S758_comb CY58_1_CD45_B02_S974_comb
C9orf152                       .                         .
RPS11                     9.2172                    8.3745
> myproject@active.assay
[1] "RNA"
```

2) at least one dimensionality reduction should exist for each cell, e.g. the tSNE coordinates.

```
> myproject@reductions$tsne
A dimensional reduction object with key tSNE_
 Number of dimensions: 2
 Projected dimensional reduction calculated:  FALSE
 Jackstraw run: FALSE
 Computed using assay: RNA

> myproject@reductions$tsne@cell.embeddings[1:2,1:2]
                             tSNE_1    tSNE_2
Cy72_CD45_H02_S758_comb    56.97119 -63.90449
CY58_1_CD45_B02_S974_comb -50.97466  38.65726
```

3) The cell identity has to be defined for each cell.

```
> myproject@active.ident[1:2]
  Cy72_CD45_H02_S758_comb CY58_1_CD45_B02_S974_comb
                        B                         T
Levels: B CAF Endothelial Macrophage NK T undefined
```

4) proper metadata has to be defined in the meta.data slot.

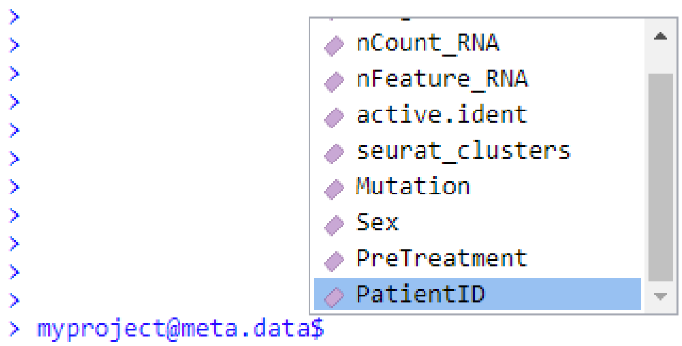


5) Three data frames are required in the obj@misc slot: obj@misc$DE where both top10 and top30 should be defined; where the basic information about the object is defined to be shown in Summary tab; and obj@misc$DataSegregation which defines which variables in the obj@meta.data slot should be used for partitioning data (more details follow in subsequent sections).

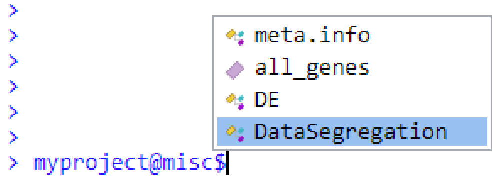


SCV enables users to work with datasets that combines multiple patients, treatments, and any other backgrounds together and be able to filter and segregate data to gain biological insights. The slot obj@meta.data is very important and any clinical information stored here can potentially be used to perform advanced data partitioning. For details please refer to section 4.7 for details.

#### 3. Creating SCV objects

To obtain a compatible rds file, the standard approach is to start with an expression matrix with cell names as column names and gene names as row names. Users may obtain the matrix in different format such as csv format, or mtx format from a Cell Ranger pipeline (10xgenomics®). The Seurat analytical pipeline takes several standard steps such as normalizing the data, finding variable features, performing PCA analysis, and finally running clustering analysis and non-linear dimensionality reduction (https://satijalab.org). To reduce un-necessary heavy computing on the front-end, SCV also required users to perform extra differential gene analysis for each cluster and store the differential gene analytical table to the Seurat object. For example the top10 gene table needs to be stored at seurat.object@misc$DE$top10 and similarly the top30 gene table needs to be stored at seurat.object@misc$DE$top30. Essential metadata is also stored on the object as a list at seurat.object@misc$meta.info. The Seurat object with this required data and analytical results is then saved as a rds file using the R built-in command saveRDS and can be then loaded by the SCV application for visualization and further analysis.

Alternatively, users can also take advantage of pre-analyzed single cell RNA-seq datasets from publications where the key information can be downloaded. We provide example script that take the pre-analyzed data and results from the Tirosh et al. 2016 datasets and processes it into an SCV-compatible Seurat v3 object. Essentially four pieces of information are required to create the previously described five slots Seurat object: 1) an expression matrix in csv format, with row names being gene names and column names being cell names; 2) a table in csv format contains the tSNE coordinates with column names being "TSNE_1" and "TSNE_2". The row names should be cell names that match the column names from the expression matrix; 3) a table containing the cluster names as the columns having row names as cell names that match the column names in the expression matrix; and 4) related metadata information in a csv format that describes the data and experiments.

A)

| | Description |
|---|---|
| Title | Dissecting the multicellular ecosystem of r |
| Authors | Tirosh I, Izar B, Prakadan SM, Wadsworth |
| Publication | Science(2016) |
| Summary | To explore the distinct genotypic and pher |
| Sample Name | Tirosh-Melanoma-Science2016-CD45P |
| Disease Type | MM |
| Tissue | Multiple |
| Enrichment | FACS CD45+ |
| Number of Cells | 2478 |
| Platform | SMARTSeq2 |
| Data Source | portals.broadinstitute.org/single_cell |
| Data POI | Tirosh[] |
| Analyst | Tirosh[] |
| Species | Homo Sapiens |
| Reference Genome | hg19 |
| ENSEMBL | NA |
| Project ID | P-20180907-0001 |
| Run ID | NA |
| Sample ID | NA |

B)

```
# Load object
myproject <- readRDS(infile)

# add metadata
meta.info.tmp <- read.csv(csvfile,
                          check.names = F,
                          row.names=1,
                          na.strings = F,
                          stringsAsFactors = F)
meta.info<-as.list(subset(meta.info.tmp, select="Description",drop=T))
names(meta.info) <- row.names(meta.info.tmp)
myproject@misc$meta.info <- meta.info

# Save object
saveRDS(myproject, file=outfile)
```

**Supplementary Figure 3.1.** Example csv file and the R command to read and load metadata to object.


#### 5. Supplementary Script

```
#!/bin/env Rscript

# Shuoguo Wang
# 2019/05/02
```

```
# Copyright: Bristol Myers Squibb


# Source Data Files:
#   - melanoma_expression.txt
#   - melanoma_cluster_assignment_portal.txt
#   - melanoma_coordinates_portal.txt
#
# Can all be downloaded from Broad Institute:
#               https://portals.broadinstitute.org/single_cell/study/SCP11/melanoma-intra-tumor-heterogeneity


### -------- functions --------- ###

replace.minus <- function(input_list) {
   input_list.collapse <- paste(unlist(input_list), collapse = "|")
   input_list.collapse.corr <- gsub("-", replacement = "_", x = input_list.collapse)
   input_list.corr <- strsplit(x = input_list.collapse.corr,split = "[|]")
   unlist(input_list.corr)
}



### ---------- setup ----------- ###

library(Seurat)
library(plyr)
library(dplyr)
data_loc   <- "./data/"
output_loc <- "./"

if(packageVersion("Seurat") < "3.0.0") { stop("Need to upgrade Seurat to >= version 3.0.0") }


### --------- pipeline --------- ###


## Load normalized data ##
df.data.temp  <- read.csv(paste0(data_loc, "melanoma_expression.txt"),
                     row.names = 1,
                     header = T,
                     check.names = F,
                     sep = "\t")
```

```
## Load classification ##
df.ident.temp <- read.csv(paste0(data_loc, "melanoma_cluster_assignment_portal.txt"),
                    row.names = 1,
                    skip = 1,
                    col.names = c("Name","Cluster","Sub_Cluster"),
                    check.names = F,
                    header = T,
                    sep = "\t")
## Load coordinates ##
df.tsne      <- read.csv(paste0(data_loc, "melanoma_coordinates_portal.txt"),
                    row.names = 1,
                    skip = 1,
                    col.names = c("Name","tSNE_1","tSNE_2"),
                    check.names = F,
                    header = T,
                    sep = "\t")

## Subset immune cells using TSNE cell names
cells <- row.names(df.tsne)
idx <- (row.names(df.ident.temp) %in% cells)
df.ident <- df.ident.temp[idx,]
idx <- names(df.data.temp) %in% cells
df.data <- df.data.temp[,idx]

## Fix cell_names which has "-"
row.names(df.ident) <- replace.minus(row.names(df.ident))
names(df.data)  <- replace.minus(names(df.data))
row.names(df.tsne)    <- replace.minus(row.names(df.tsne))

# Create seurat 3 object
myproject <- CreateSeuratObject(counts = df.data)
myproject@active.assay <- "RNA"

## Add meta.info
meta.info.tmp <- read.csv(paste0(data_loc, "Tirosh-MM-2016-CD45P.csv"),
                    check.names = F,
                    row.names = 1,
                    na.strings = F,
                    stringsAsFactors = F)
meta.info <- as.list(subset(meta.info.tmp, select = "Description", drop = T))
names(meta.info) <- row.names(meta.info.tmp)
myproject@misc$meta.info <- meta.info

## set cell ident
```

```
cell_ident <- droplevels(df.ident$Cluster)
names(cell_ident) <- rownames(df.ident)
myproject[["Tirosh.ident"]] <- cell_ident
Idents(myproject) <- myproject[["Tirosh.ident"]]

## set dr
dr_tsne <- CreateDimReducObject(embeddings = as.matrix(df.tsne),assay = "RNA",key = "tSNE_")
myproject[['tsne']] <- dr_tsne

## compute DEG
myproject@misc$all_genes <- rownames(myproject)
myproject@misc$DE$all <- FindAllMarkers(object = myproject,
                                        only.pos = FALSE,
                                        min.pct = 0.05)
myproject@misc$DE$all %>% group_by(cluster) %>% top_n(10, avg_logFC) %>% as.data.frame()
-> myproject@misc$DE$top10
myproject@misc$DE$all %>% group_by(cluster) %>% top_n(30, avg_logFC) %>% as.data.frame()
-> myproject@misc$DE$top30

## add meta
## The following meta.data was manually curirated from Tirosh supplementary table S1 and S2:
sample_id <- table(tolower(unlist(lapply(colnames(myproject),
                                  function(x) {substr(x,1,4)[[1]][[1]]}))))
sample_list <- tolower(unlist(lapply(colnames(myproject),
                              function(x) {substr(x,1,4)[[1]][[1]]})))
PatientID <- mapvalues(sample_list,
                       from = names(sample_id),
                       to =
c("Mel53","Mel58","Mel60","Mel72","Mel74","Mel79","Mel80","Mel81","Mel84","Mel88","Me
l89","Mel94"))
Mutation <- mapvalues(sample_list,
                      from = names(sample_id),
                      to = c("Wild-type","Wild-type","BRAF","NRAS","na","Wild-
type","NRAS","BRAF","Wild-type","NRAS","na","Wild-type"))
Sex <- mapvalues(sample_list,
                 from = names(sample_id),
                 to   = c("F","F","M","F","M","M","F","F","M","M","M","F"))
PreTreatment <- mapvalues(sample_list,
                          from = names(sample_id),
                          to = c("N","Y","Y","Y","Y","N","N","N","N","Y","N","Y"))
names(PatientID) <- colnames(myproject)
names(Mutation) <- colnames(myproject)
names(Sex) <- colnames(myproject)
names(PreTreatment) <- colnames(myproject)
```

```
myproject@meta.data$Mutation <- Mutation
myproject@meta.data$Sex <- Sex
myproject@meta.data$PreTreatment <- PreTreatment
myproject@meta.data$PatientID <- PatientID

myproject@misc$DataSegregation <- list("Mutation" = names(table(myproject@meta.data$Mutation)),
                       "Sex" = names(table(myproject@meta.data$Sex)),
                       "PatientID" = names(table(myproject@meta.data$PatientID)),
                       "PreTreatment" = names(table(myproject@meta.data$PreTreatment)))

saveRDS(myproject, paste0(output_loc, "tirosh_seurat3.RDS"))
message('Finished, file output to: ', paste0(output_loc, "tirosh_seurat3.RDS"))
```
